# Seq2KING: An unsupervised internal transformer representation of global human heritages

**DOI:** 10.1101/2025.06.17.660172

**Authors:** Bhavana Jonnalagadda, eMalick Njie

## Abstract

Determining the intricate tapestry of human genetic relationships is a central challenge in population genetics and precision medicine. We propose that the principles of lexical connectivity, which words derive meaning from their contextual interactions, can be adapted to genetic data, enabling transformer models to reveal that individuals with higher genetic similarity form stronger latent connections. We explored this by transposing KING kinship-related matrices into the (query, key, value) QKV latent space within transformer models and determined that attention mechanisms can capture genetic relatedness in an unsupervised fashion. We found that individuals had an attention weight connectivity of 85.34% (p<0.05) if they were from within the same continent, compared to if they were from other continents. Surprisingly, we found that some encoder layers required inversion of their latent representations for this connectivity to become obvious. Lastly, we used BERTViz to create human-readable hyper-dense connectivity patterns among individuals. Our approach is purely based on attention, which yields a non-discrete spectrum of relatedness, and thus uncovers patterns on first principles. Seq2KING addresses the significant challenge of discovering population structures to construct a global human relatedness map, without relying on predefined labels. Our excavation into the latent space is a paradigm shift from legacy-supervised genetic methodologies, which presents a new way to understand the human pangenome as well as discern population substructures for creating precision genetic medicines.

**Non-Expert Description:** Is it possible to build artificial intelligence (AI) to read the human genome as a first language? Why would one want such AI? We at Ecotone believe that such AI will provide the genetic coordinates needed to manufacture CRISPRs medications to cure about ∼10,000 genetic diseases. How does one build such AI? Our recently released model dnaSORA proposed a means to assign meaning to every single token (typically referred to as a base) of all 3 billion tokens in the human genome (Koreniuk & Njie, 2025). This builds the vocabulary for reading the human genome as a first language. For dnaSORA to work, it needs to know the heritages of people that are in its model of our genetics. We mostly rely on country, culture and geography to determine our heritages, but this is too error-prone for dnaSORA. Also error-prone in our experience are legacy genetic approaches such as those used by 23andMe.

Our research here introduces Seq2KING, a new artificial intelligence method that is based on excavating the insides of transformers to uncover hidden patterns of genetic relatedness among people around the world—without needing any prior labels or categorizations. The key innovation of Seq2KING is applying the principles of lexical connectivity— how words derive meaning through their relationships to other words—to genetic data. Just as “dog” gains meaning through its connections to words like “pet,” “animal,” and “loyal,” we show that individuals’ genomes can be understood through their genetic connections to others. We start by converting raw genetic data into a compact kinship matrix (using a tool called KING) that summarizes how closely everyone is related. We then feed these kinship values into a transformer model—the same kind of AI behind cutting-edge language tools like ChatGPT.

Inside the transformer, special components called “attention heads” learn which individuals are most similar, strengthening links between people from the same region and showing subtler connections across continents. Unlike legacy approaches that rely on discrete pre-defined categories, Seq2KING provides continuous measures of relatedness, allowing us to visualize connections between any individual and all other humans. Additionally, because Seq2KING operates directly within the transformer’s internal reasoning system, it can be seamlessly integrated as a component within larger genome interpretation systems—essentially functioning like high-speed cache memory for heritage assignments, dramatically improving both efficiency and scalability. By examining these attention patterns, we can reconstruct familiar population groupings—such as European, African, and Asian heritage—entirely by the model’s internal logic. Finally, we use a visualization technique (BERTViz) to turn these dense connection maps into intuitive diagrams that highlight population connections between individuals.

Because our approach doesn’t rely on pre-assigned labels, it offers a truly unbiased way to explore human population structure. This could help scientists trace migration routes that resulted in the peopling of the continents, find subtle subgroups within larger populations, and remove “background noise” in genetic studies of disease. Ultimately, Seq2KING paves the way for more precise genetic maps of all humans, revealing the natural “family trees” hidden in our DNA and bringing us one step closer to reading the human genome as a first language.

## 1 Introduction

Understanding the genetic relatedness of any given human to all humans (∼8 billion pangenome) — is an open and fundamental question. Answering this question gives insight into the peopling of the continents and origins of the ∼950 human heritages in existence today (UNESCO World Heritage Centre, 2025). Moreover, the nuances of heritage are a major source of noise for efforts to identify sections of the genome that cause genetic diseases when altered. Thus, having a high-resolution relatedness map of all humans enables the removal of this noise, which can directly lead to next-generation medicines that are reliant on genetic precision such as CRISPR.

Advancements in whole-genome sequencing have led to the rapid growth of pangenome collections with consistent increases in the breadth of genetic representations of human heritages (Liao *et al*., 2023). Thus, one can envision a connectivity model where a single individual is genetically linked to all other humans.

Although parallels between language space and heritage spaces are conceptually compelling, the application of lexical connectivity to genetic data remains an uncharted frontier. Our work seeks to explore these relationships using latent internal representations in attention-based transformer architectures. Such architectures have revolutionized natural language processing by capturing ultra-fine-grained lexical meanings in the language space. An example is shown in Figure 1, where meaning within the attention layers of a deep learning (self-attention transformer) model is contextual based on the input. Here ‘man’ has high pronoun attention in a sentence with ‘he’ and not in the same sentence whose only difference is ‘she’. We extended these methods to decode the relationships in the heritage space.

**Figure 1:**
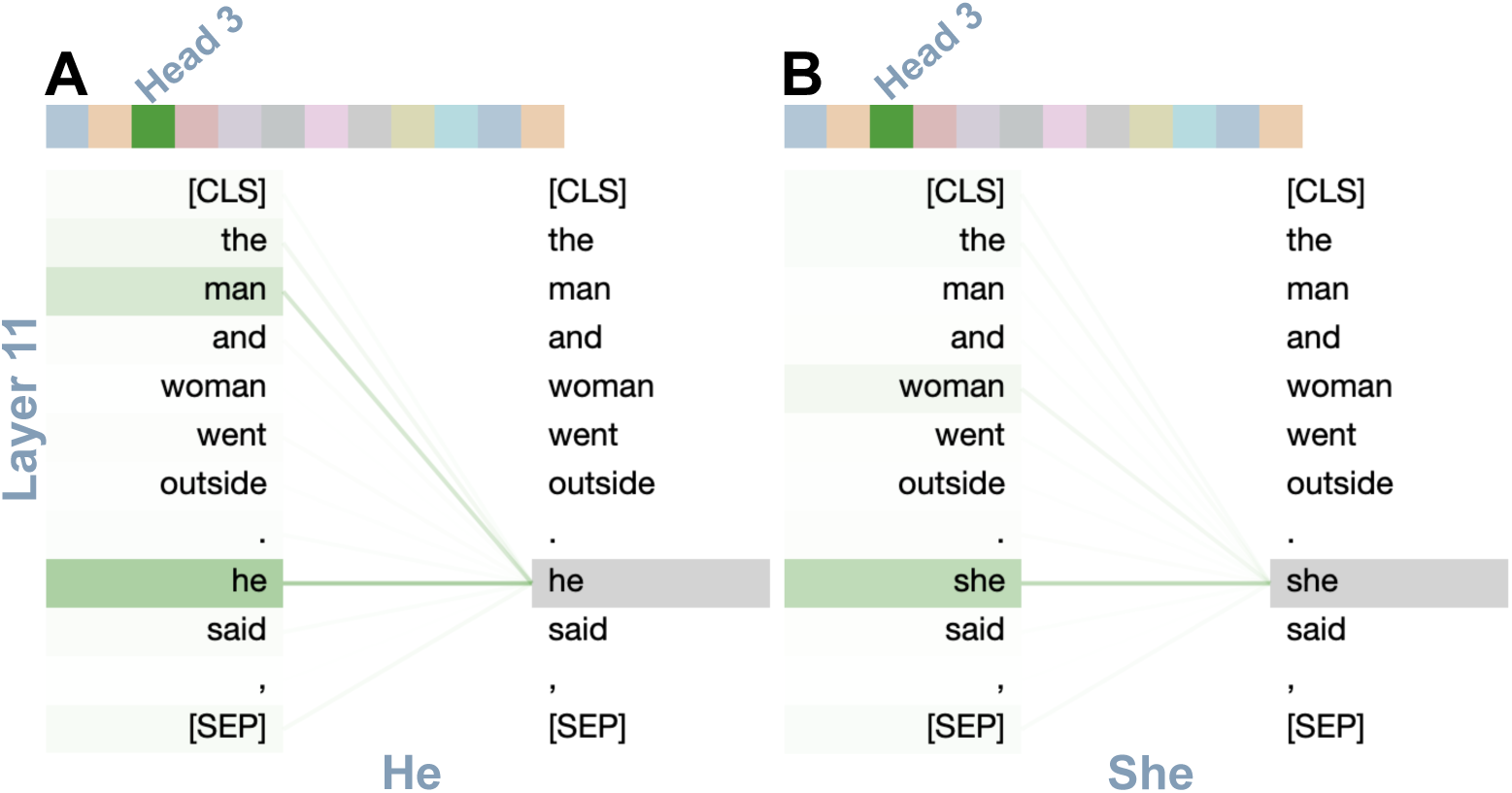
An example of the visualization of the attention weight strengths of an encoder layer (layer 11), of the base Google BERT (case-agnostic) model (Devlin *et al*., 2018), for head 3; the chosen input and weights highlight gendered attention. **A**: The second sentence starting with “he”. **B**: The second sentence starting with “she”. Note attention (green line) to ‘man’ only occurs in A as there is lexical masculine connectivity of ‘he’ to ‘man’. Made with BERTViz (Vig, 2019).

### 1.1 Contributions

We introduce Seq2KING, a novel approach that utilizes a self-attention transformer architecture designed specifically for genetic relationship analysis. Its architecture is inspired by Seq2Seq (Sutskever *et al*., 2014), where the input language sequence is processed into a large fixed-dimensional vector representation (a context vector) and then decoded into another language. Our model translates from an input sequence, but instead of resulting in an equivalent output sequence, it converts to KING values (elaborated below). Our approach addressed several challenges.

First, the raw 30x coverage ATGC whole-genome representation for a typical human is approximately 50 gigabytes. The total representation for the current eight billion humans is ∼400,000 petabytes. Such pangenome representations cannot be computed or stored in memory using advanced hardware. We side-step this issue by employing dimensionally compressed KING matrices, thus theoretically enabling the representation of all humans.

Second, we excavated the QKV’s latent space. This latent space is a challenging non-linear environment to explore, but is the most granular information of artificial intelligence and has been successfully explored recently to reveal the polysemanticity of neural nets to explain AI (Ameisen *et al*., 2025; Lindsey *et al*., 2025). In an unsupervised fashion, we found that attention weight connectivity was strong within the latent space for individuals from the same continent and weak for those not from the same continent.

Finally, we anticipate the difficulty of visualizing and interpreting QKV attention-weight level data graphs of the relatedness of one human to all other humans. Thus, we introduce the use of BERTViz in genetics to enable sensible visualization and fast human-readable analyses of this high-density data.

These insights are collectively Seq2KING which is a paradigm shift from legacy methods and is the first demonstration of the potential of latent space excavation (a.k.a., neural excavation, see Related Work) in genetics and medicine. We foresee its generalization for understanding the forking of DNA as humans peopled the continents, forming today’s ∼950 heritages. It is also useful for the population substructure assignments required for the creation of precision genetic medicines.

## 2 Related Work

### 2.1 Legacy approaches to population inference

Population inference, a key genetic task, involves understanding the structure of populations based on genetic variation. Traditionally, non-deep learning methods such as maximum likelihood estimation have been widely used (Tang *et al*., 2005). Another approach uses Bayesian models to manage the complexity of large datasets, which can include millions of individuals and genetic markers (Gopalan *et al*., 2016). Bayesian methods are effective for estimating population structures; however, they face limitations in terms of computational scalability and the complexity of nonlinear data patterns. Evaluations comparing legacy and newer nonlinear techniques, such as t-SNE and UMAP, have shown that graph-based algorithms outperform principal component analysis (PCA) in detecting intricate population structures (Njie, 2018; Ubbens *et al*., 2022).

ADMIXTURE is another widely used approach in population inference, that leverages model-based estimation of genetic ancestry. The original ADMIXTURE model, developed by Alexander and colleagues, combined PCA and constraint based model searches to optimize ancestry estimates (Alexander *et al*., 2009). More recent adaptations have enhanced the efficiency with likelihood-free parameter initialization, as seen in a faster version of ADMIXTURE (Cabreros & Storey, 2019; Santander *et al*., 2024).

### 2.2 Deep learning techniques in genetics and population inference

Despite the success of legacy methods such as ADMIXTURE, these techniques face challenges in processing largescale genetic data due to the increasing use of whole genome sequencing, now encompassing millions of single nucleotide polymorphisms (SNPs). Deep learning-based approaches are emerging as promising alternatives owing to their scalability and ability to capture nonlinear patterns in data. Neural networks, particularly autoencoders, have been applied to replicate ADMIXTURE, but have not yet surpassed legacy methods in population inference tasks (Mantes *et al*., 2022). Recent work using transformer networks, but not specifically focused on population inference, has demonstrated the viability of applying these advanced architectures in genetics, particularly for genomic context identification and molecular phenotype prediction (Brixi *et al*., 2025; Dalla-Torre *et al*., 2023; Nguyen *et al*., 2023). However, the transformer methods used in these studies have not explored the use of the latent space inside attention-based transformers, a promising area of research and direction. Legacy methods have also been limited in their ability to perform exploratory data analysis, particularly for visualizing and interpreting relationships between individuals in large-scale genetic datasets.

### 2.3 KING kinship relatedness

Transforming genetic data, such as SNPs, into KING kinship relationships facilitates faster and more accurate relationship analysis of genetic data. KING is a relationship-inference algorithm for capturing unknown population substructures (Manichaikul *et al*., 2010). Briefly, it uses identical-by-descent (IBD) statistics between each pair of individuals, modeling the genetic distance between a pair of individuals as a function of their allele frequencies and kinship coefficient. KING outputs kinship coefficient matrices that are independent of sample composition or population structure. This mechanism makes KING particularly valuable for analyzing the global pangenome, where genetic diversity and substructure present unique challenges. Moreover, KING output is particularly desirable because it can be derived from petabyte quantities of whole genomes and remarkably flattens this to megabyte size data. This flattened data is a manifold compression that contains essential relatedness information from billions of ATGC raw genomic elements. The manifold is a high-level feature space composed of thousands of dimensions that cannot be interpreted by the human eye. We have previously demonstrated unsupervised dimensionality reduction techniques on KING matrices that have successfully revealed global population structures (Njie, 2018). However, fully leveraging the KING manifold remains an unmet challenge.

### 2.4 Self-attention transformers

Deep learning techniques have become increasingly prominent in various scientific fields, including genetics, owing to their ability to model complex, nonlinear relationships. Neural networks, in general, have revolutionized data processing, but the transformer model stands out because of its capability to manage long-range dependencies and parallelize computations. The attention-based transformer architecture, first introduced by Vaswani et al., has been adapted across many domains beyond genetics, such as natural language processing and computer vision, because of its ability to dynamically focus on relevant inputted data (Kaiser *et al*., 2017; Vaswani *et al*., 2023).

The success of the transformer architecture lies in its attention mechanism, which allows the model to weigh the importance of different inputted elements differently, enabling it to capture complex relationships and dependencies in the data. This capability has been particularly evident in the development of large language models such as ChatGPT and Claude, which have demonstrated unprecedented abilities in understanding and generating human language. The success of these models in capturing complex patterns and relationships in sequential data suggests similar potential in genetic analysis, where understanding relationships between genetic markers and individuals is crucial.

### 2.5 Neural excavation

Neural excavation is an emerging field of artificial intelligence that refers to exploring internal representations inside AI models for utility and insight. For instance, recent work by Anthropic investigated whether objects and concepts are represented in individual neurons or as distributed features across neurons (Ameisen *et al*., 2025; Lindsey *et al*., 2025). QKV values of transformers compose a latent representation—a compressed mathematical encoding that captures real-world information in abstract numerical structures.

DeepSeek demonstrated that the computational overhead of attention mechanisms can be dramatically reduced through Multi-Head Latent Attention (MLA), which compresses Key-Value caches into secondary latent representations before expanding them back for computation. This innovation reduced the key-value cache by a factor of 57 and enabled text generation more than six times faster than traditional transformers (DeepSeek-AI *et al*., 2025). The successes of Anthropic and DeepSeek encourage further exploration of latent spaces.

As we previously discussed, to generate realistic videos, world models such as SORA are likely to encode fundamental functions of nature —motion, gravity, physics —-within their latent representations (Koreniuk & Njie, 2025). We term the emerging discipline of building tools to unearth such encoded functions ‘neural excavation’. Importantly, these functions are not limited to those already known to science, making this an exciting frontier. In genetics, the fundamental functions governing genomic relationships remain largely unknown, making Seq2KING’s exploration of transformer latent space particularly valuable.

## 3 Results

### 3.1 Model architecture

Traversing the spectrum of genetic similarity, we hypothesized that a transformers attention mechanism will reinforce bonds among individuals within the same population while sustaining subtle, organized interactions across different populations. We tested this hypothesis by building Seq2KING, a new, unsupervised approach for understanding genetic relatedness.

To begin investigating, we tested various architectures and hyperparameters of transformers. The top combinations (as determined by loss) and their hyperparameters are displayed in Table S1. The chosen architecture for Seq2KING after testing is displayed in Figure 2, with its hyperparameters indicated in the bolded row of Table S1. Final training and validation MSE loss were performant, at 3.1*e*^−5^and 2.3*e*^−5^ respectively.

**Figure 2:**
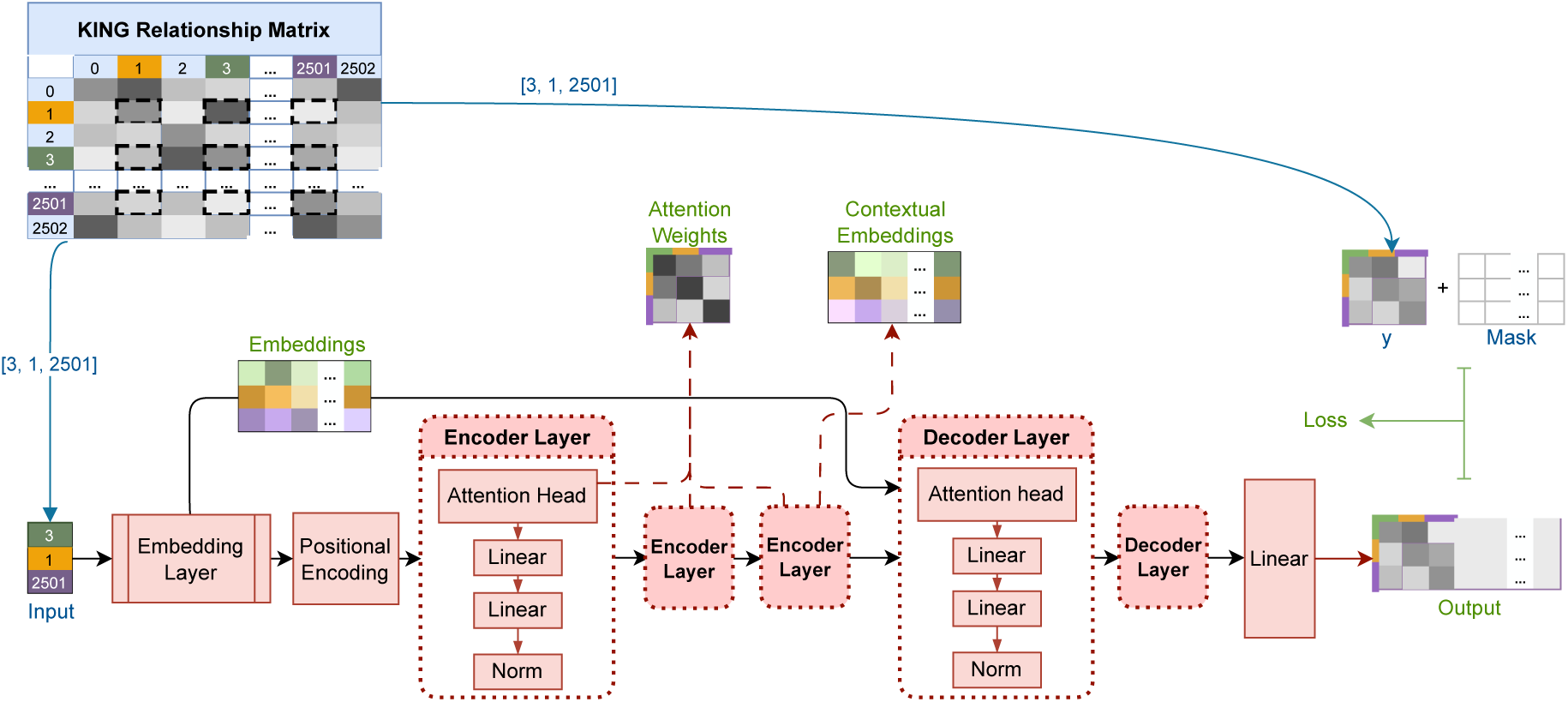
The final transformer neural network architecture used in Seq2KING, with the data flow also illustrated. The given input is a list of tokens representing the *n* indices of given individuals (in this example, [3, 2, 2501]) and the output is compared against the given KING values taken from the reflected KING matrix. The blue information is related to the original KING matrix that was inputted, the green information is data we inspect in order to analyze the model, and the red is the model architecture itself. See Figure 8 for details about the reflected filled matrix.

We generated a KING matrix extracted from whole genomes of 2,503 individuals from the 1000 Genomes Project as previously described (Njie, 2018). The information space of all chromosomes in whole genomes challenge modern hardware, therefore we extricated chromosome 1. We’ve shown previously this chromosome is sufficient to capture global substructure (Njie, 2018). The matrix itself is composed of values that are filled above the diagonal and empty under the diagonal. We performed cluster analyses using UMAP of this version of the matrix and a second version where we reflected values above the diagonal to the empty space below the diagonal. We found the reflected version to result in clustering of related populations best (Figure 8). We therefore trained our models on the reflected dataset. Population labels were excluded in training (i.e., unsupervised).

Our model processes input as a list of tokens representing the *n* indices of given individuals. The output consists of a *n* × *n* matrix predicting the KING relations between the *n* individuals. This produces a latent space of learned embeddings that is a high-dimensional representation of genetic relationships.

### 3.2 Multiple heads in encoder layer

We experimented with multiple attention heads in the encoder layer. Increasing the number of heads did not provide additional benefits. The attention patterns exhibited lower correlation across heads, leading to less coherent representations. Furthermore, models with multiple heads achieved similar final loss values compared to those with a single attention head. This suggests no advantage in increasing model complexity in this regard. We also observed that additional encoder and decoder layers did not improve performance.

### 3.3 UMAP embeddings

We extracted learned embeddings and visualized their structure using UMAP. Despite never explicitly receiving population labels, the embeddings grouped into distinct populations (Figure 3). This population structure was our first qualitative indication that our model learned genetic relationship patterns without supervision.

**Figure 3:**
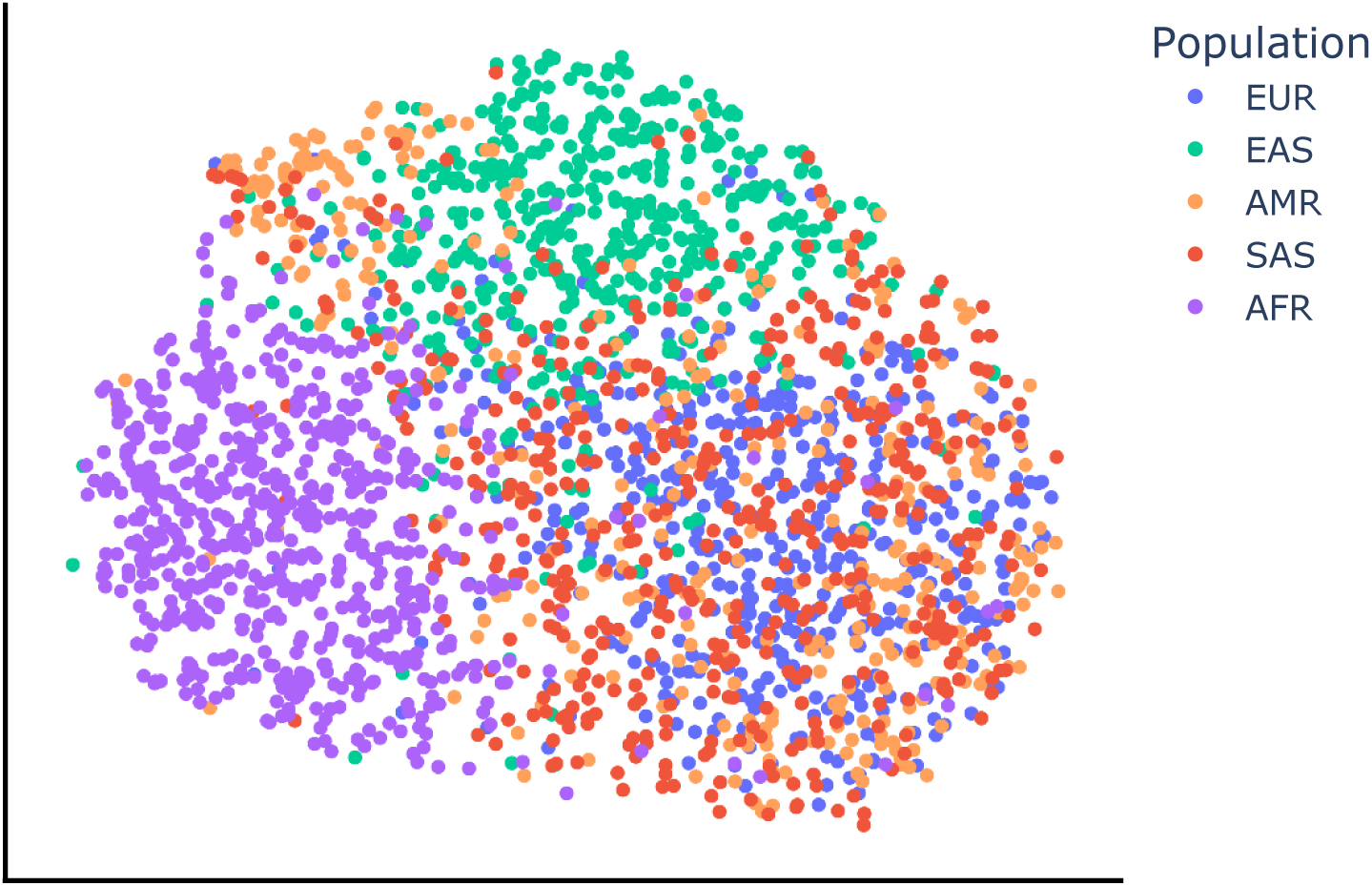
Dimensionality reduction using UMAP to 2D of the embeddings taken from the embedding layer of Seq2KING, colored by continental population label (unseen and entirely learned by model).

### 3.4 Attention within encoder layers

#### 3.4.1 Visualization of attention

In the following, continental acronyms from the 1000 Genomes Project are: EUR - Europe, EAS - East Asia and Pacific, AMR - Americas, SAS - South Asia, AFR - Africa. We examined the attention mechanisms within the encoder layers to understand how the model distributed focus among individuals. We anticipated geographically co-localized individuals to have higher relatedness and higher QKV weights than geographically distant individuals. For example, because the Americas are majority composed of European decent compared to Asia, individuals with the EUR population label should show more connectivity to AMR individuals than to SAS individuals, and the most connectivity to other EUR individuals.

BERTViz is a tool for visualizing attention weights (Vig, 2019). Its visualizations are scalable with comparisons of a few weights being as intuitive as comparisons of thousands of weights. We therefore used BERTViz to display attention-based connectivity. We also use heatmaps of the same attention values to supplement BERTViz.

The visualized attention weights are shown for three selected groups of individuals. The first group is AMR and EAS population individuals, the second of AFR and SAS individuals, and the last group is of a randomly selected set of individuals that represent every continental population in the dataset (collectively referred to as the Three-Sample Subsets).

#### 3.4.1 Positive correlation in encoder layer 1

We display attention weights of all encoder layers in Figure 4. We note that in layer 1, we see positively correlated attention weights of individuals to themselves, aka strong self-attention (strength discussed below with results from the other layers. Quantification in Table 1 and Table 2). The positively correlated attention weights agree with our hypothesis that the attention mechanism will reinforce bonds among individuals within the same population.

**Figure 4:**
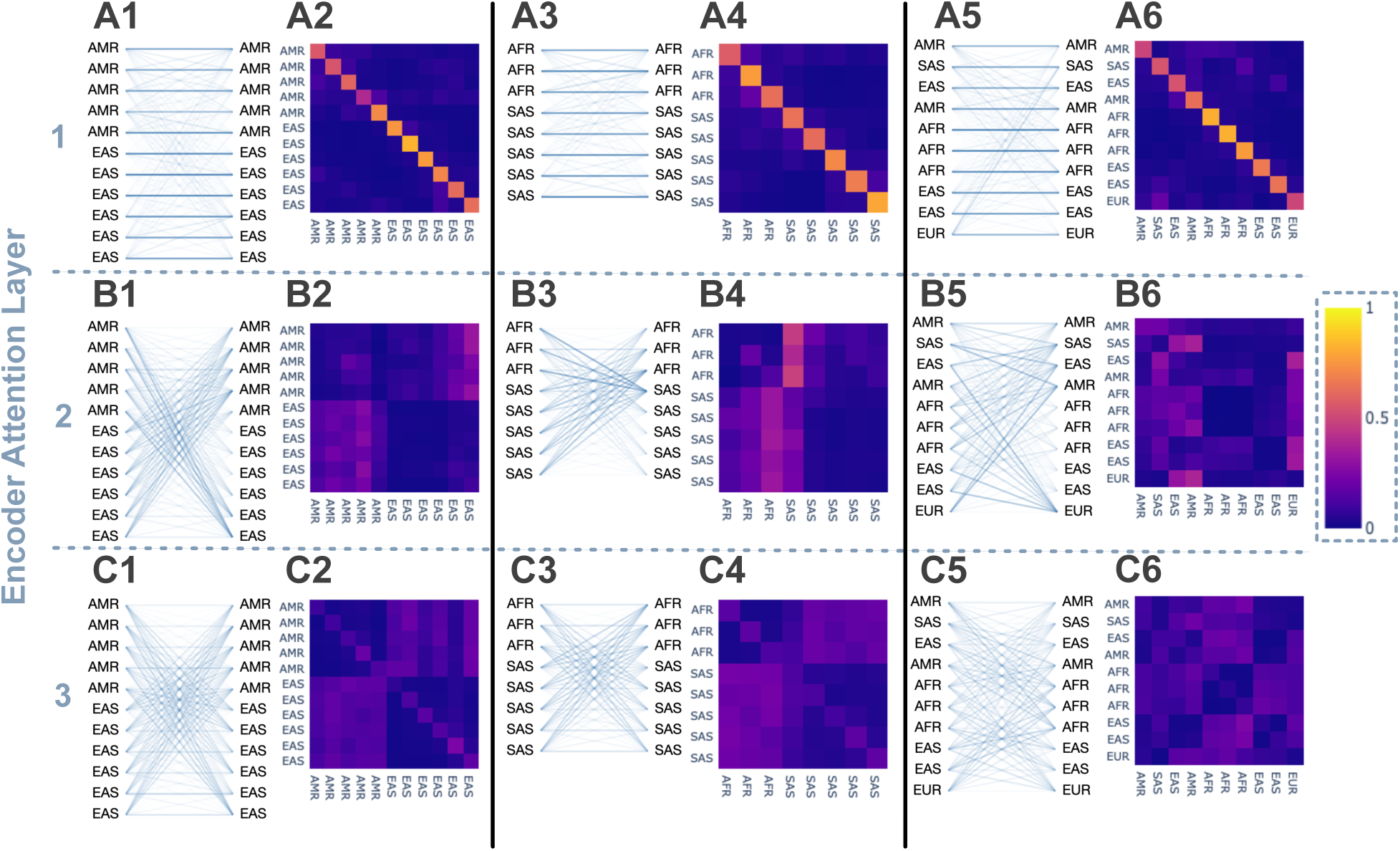
The attention weights of each encoder layer, illustrated through BERTViz and heatmaps, for selected sets of individuals. **A**: Layer 1. **B**: Layer 2. **C**: Layer 3. **Odd Numbers**: BERTViz of the weights. **Even Numbers**: Same information displayed in a heatmap. **1-2**: Selected individuals to display relation between AMR and EAS populations. **3-4**: Selected individuals to display relation between AFR and SAS populations. **5-6**: A random selection of individuals that include all continental populations. Note the anti-correlation in layers 2 and 3.

**Table 1:**
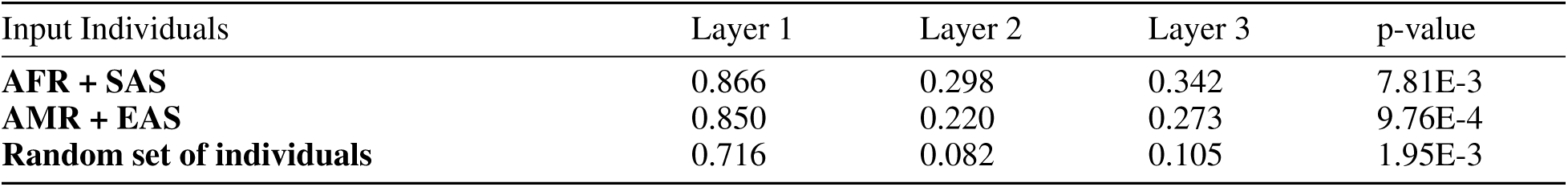
The average intra-connectivity ratios per layer, and their associated p-values, for the three cohorts of sample individuals.

**Table 2:** The average intra-connectivity across all individuals in the original dataset, averaged also across populations, per layer, along with associated p-value. Layers 2 and 3 have inverted values.

| Layer | Average intra-connectivity ratio | p-value |
| --- | --- | --- |
| **1** | 0.743 | 0.031 |
| **2 (inverted)** | 0.853 | 0.031 |
| **3 (inverted)** | 0.801 | 0.031 |

#### 3.4.2 Negative correlation in encoder layers 2 and 3

In contrast to the first encoder layer, layers 2 and 3 exhibited negative correlation in attention weights (Figure 4). This anti-correlation was observed on all runs of the final architecture of the model. We therefore inverted these attention weights to convert these values to high positive correlations for individuals of the same population. An example of this inversion is shown in Figure 5 for AFR-SAS individuals. Noted thickened lines are due to the original values being close to 0 and the inverted values being close to 1.

**Figure 5:**
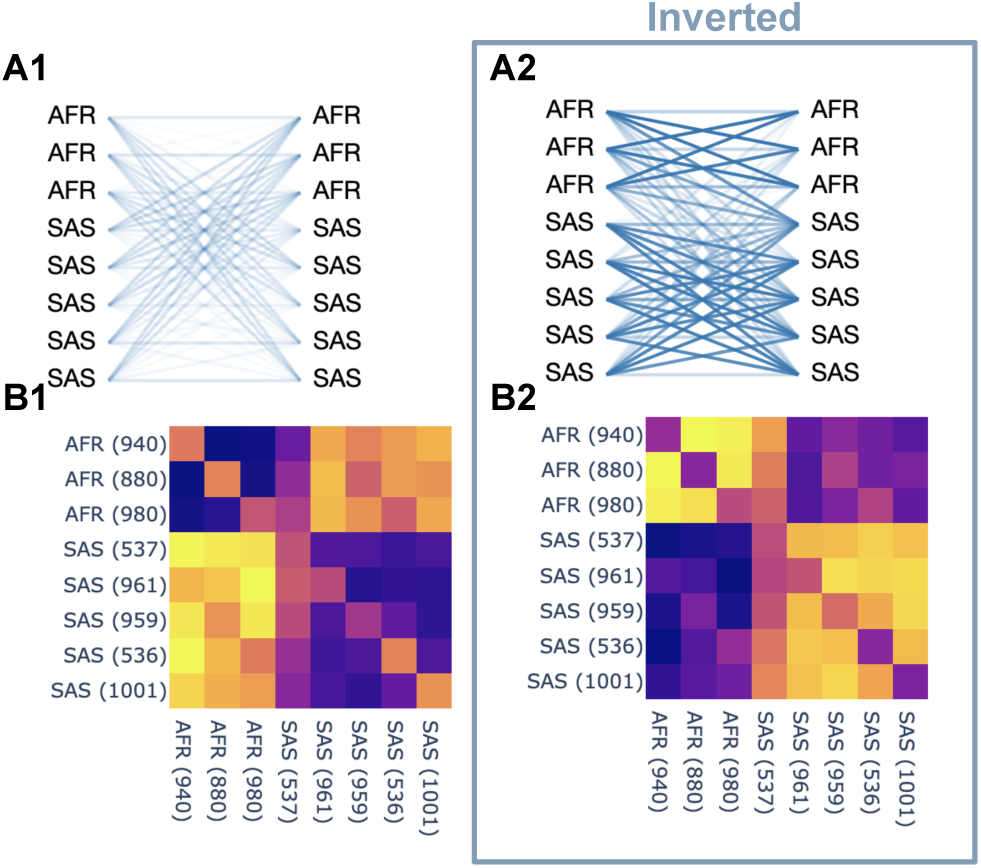
Example of numerical inversion on anti-correlated attention weights. Strong anti-correlation was observed in layers 2 and 3 (Figure 4). This is a surprising feature that we leveraged and made more human-readable by inversion. Example here from layer 3 shows the original values for AFR-SAS **(A)** and the inverted values **(B)**. Note change of line weight thickness between A and B. This is due to weights inverting from 0 towards 1. For example, given a weight of 0.02, the inverted weight is now 0.98 (given max line thickness corresponds to weight of 1.0).

To quantify these patterns, we calculated the average intra-population and inter-population attention weights across all individuals (Figure 7). We defined intra-percentage and inter-percentage as the proportion of attention weight values connecting individuals within the same population and spanning different populations respectively. We performed statistical tests (see Methods) on the Three-Sample Subsets (Figure 4, Figure 6, Table 1). Note encoder layers 2 and 3 were inverted. We found statistically significant intra-population connectivity in all layers (p<0.05).

**Figure 6:**
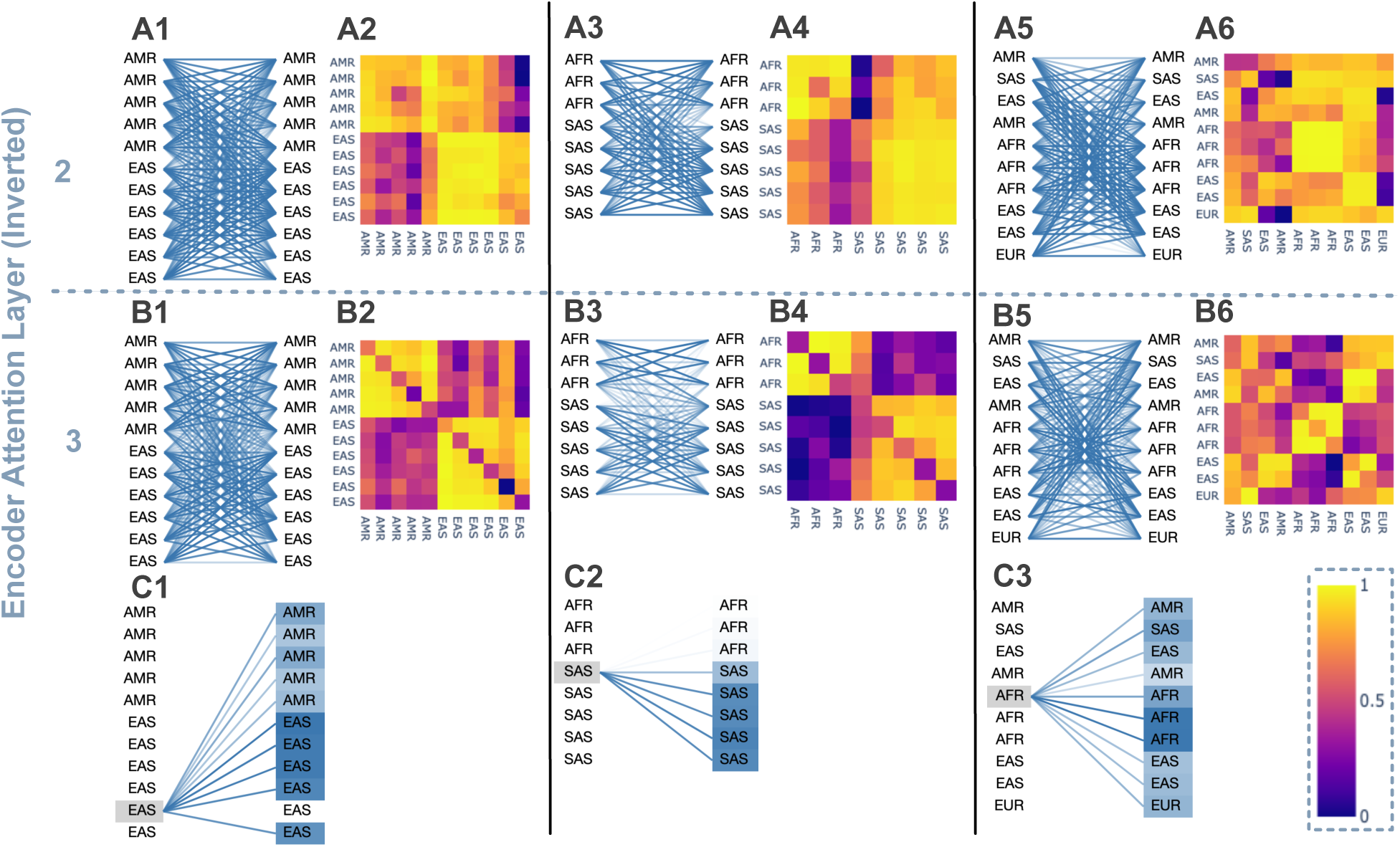
The attention weights for encoder layers 2-3, numerically inverted. **A**: Layer 2. **B**: Layer 3. **A-B Odd Numbers**: BERTViz of the weights. Note that the weights are higher numerically (“stronger” line thickness) compared to Figure 4 due to the inversion, as many weights were low value. For example, given a weight of 0.02, the inverted weight is now 0.98 (given max line thickness corresponds to weight of 1.0). **A-B Even Numbers**: Same information displayed in a heatmap. **C**: A selected individual with its weights/connections highlighted, for each selected input sets. **A-B 1-2**: Selected individuals to display relation between AMR and EAS populations. **A-B 3-4**: Selected individuals to display relation between AFR and SAS populations. **A-B 5-6**: A random selection of individuals that include all populations. GIFs demonstrating dynamic weight assignments available online at Supplemental Link Section 6.1.

**Figure 7:**
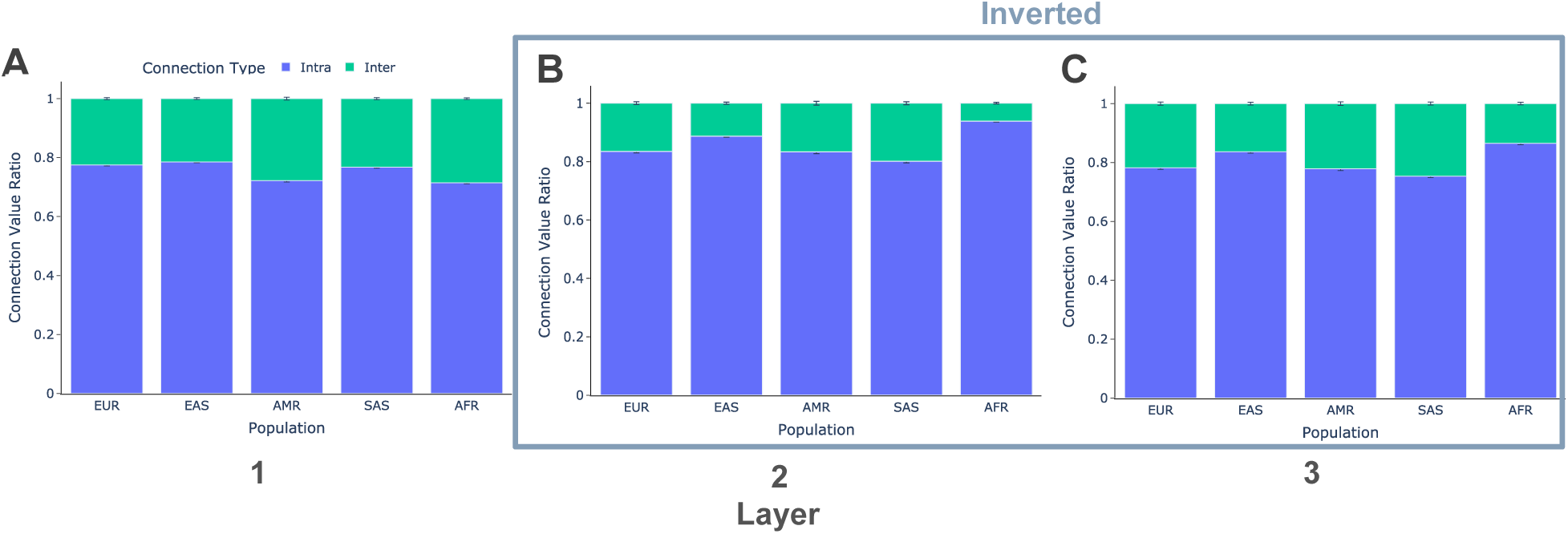
All average intra/inter percentage of weights. Intra-percentage is defined as the percentage of encoder attention weights values that connect to the same population (within-population); inter-percentage is the same for weights that connect to other populations (between-population). Layers 2 and 3 numerically inverted for visualization purposes. Note all values pass significance testing at *α* = 0.05. **A**: Layer 1 averages. **B**: Layer 2 averages, inverted. **C**: Layer 3 averages, inverted.

**Figure 8:**
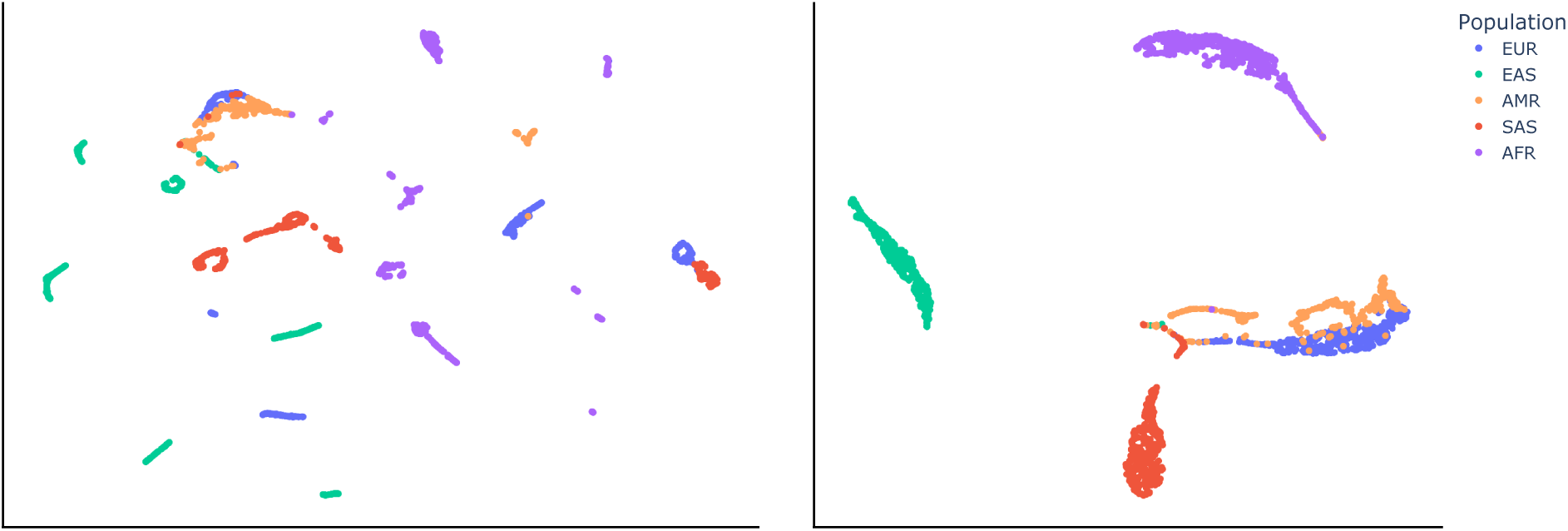
Comparison on population structure in unreflected and reflected KING matrices. KING kinship data is composed of a matrix with values only above the diagonal. **A**: UMAP of this data results in clustering that is poorly correlated to continental labels. **B**: Reflection of all values across the diagonal so the bottom of the diagonal is filled with duplicate values results in clusters that are more clearly correlated with continental labels. This “filled” matrix was therefore employed as the input for Seq2KING (see Methods). Continental labels: EUR - Europe, EAS - East Asia and Pacific, AMR - Americas, SAS - South Asia, AFR - Africa.

We also examined the connectivity ratios averaged across all individuals in Figure 7, shown in Table 2. Layer 2 average intra-connectivity was 85.34% (p<0.05).

### 3.5 Contextual embeddings

Contextual embeddings provide dynamic representations of individuals. They capture relationships within the sequence rather than as static embeddings. Unlike standard embeddings (discussed in the above section “Embeddings”), which remain fixed regardless of position, contextual embeddings adapt based on surrounding individuals and the model’s learned attention patterns. To assess the positional invariance of these contextual embeddings, we conducted experiments by shifting an individual’s position from index 0 to index 10 within a fixed sample of 10 individuals. The remaining individuals in the sequence maintained their original positions. We then computed the cosine similarity between the contextual embeddings of the given individual across different input lists. The resulting cosine similarity matrix, visualized in Figure S2, demonstrated that the contextual embeddings remained consistent regardless of positional shifts. This invariance indicates that the model learns representations independent of absolute positioning, suggesting robustness in how it captures individual relationships. However, this invariance did not hold across different combinations of individuals, limiting the ability to combine different runs to provide a full comprehensive view of all contextual embeddings — all individuals present in a combined contextual embedding matrix to be able to directly compare them.

Additionally, the contextual embeddings exhibited separation patterns corresponding to population clusters. Despite never receiving explicit population labels, individuals from the same population tended to have more similar contextual embeddings. This reinforces the notion that the transformer inferred genetic structures purely from the KING matrix. These results further highlight the model’s ability to capture meaningful genetic information in a structured manner. It does so without requiring explicit supervision or prior knowledge of population groupings.

## 4 Methods

Precise details on implementations and usage of statistical methods, built models, data, and evaluation metrics are located at Supplemental Section 6.1.

### 4.1 Data

#### 4.1.1 1000 Genomes project data

We obtained genetic variation data from the 1000 Genomes Project, a large-scale international initiative designed to catalog human genetic diversity (Auton *et al*., 2015; Fairley *et al*., 2020). The project offers a comprehensive resource that captures genomic variation across diverse global populations. We chose this dataset because it provides extensive, high-quality data that is ideal for studying population structure and kinship relationships. For our analysis, we selected 2503 individuals (representing a diverse spread of heritages) using the chromosome 1 variants derived from the GRCh37 reference genome. This provides a robust and representative sample for training and evaluating our model. Continental labels: EUR - Europe, EAS - East Asia and Pacific, AMR - Americas, SAS - South Asia, AFR - Africa. The model was trained on all 2503 individuals from these geographic locations. Note the model is non-parametric - additional new individuals require re-training.

#### 4.1.2 KING relationship inference matrix

Whole genomes are computationally expensive particularly for relatedness maps of the human pangenome. We therefore employed KING pairwise relatedness coefficients, as they enable us to condense the genomic data into a much smaller set of values, while preserving critical information about genetic similarity (Manichaikul *et al*., 2010). KING employs a fast and efficient algorithm that compares genotype data across individuals to estimate kinship coefficients. Briefly, it uses identical-by-descent (IBD) statistics between each pair of individuals, modeling genetic distance between a pair of individuals as a function of their allele frequencies and kinship coefficient. Its broad applicability has been demonstrated in various studies—from reconstructing small family pedigrees in rare disease research to defining relationships among continental populations (Conomos *et al*., 2015; Manichaikul *et al*., 2010). We previously demonstrated global population structure can be derived from KING data (Njie, 2018).

The KING matrix was imported in Python and transformed into a square upper-triangular matrix, where each entry at position (k, n) represents the genetic relatedness between individuals k and n. To ensure a symmetric representation of kinship relationships, we constructed a reflected “filled” matrix by mirroring the upper-triangular portion into the lower-triangular section. This reflected representation was tested to determine whether symmetrical kinship encoding improved model performance; the filled KING matrix was retained in the final model architecture as it was found to provide a more complete representation of genetic relationships. As demonstrated in Figure S1, the KING matrix values correlate with intra- and inter-population connectivity (but is not a perfectly linear relationship). Higher, positive KING values correspond to intra-population relationships (within a population), reflecting greater genetic similarity. Conversely, inter-population connectivity (between different populations) is characterized by lower or negative KING values, indicating weaker genetic relationships between individuals from different continental populations.

### 4.2 Network structure and evaluation metrics

We aimed to optimize the performance of our transformer-based model by systematically exploring its network architecture and hyperparameter settings. The hyperparameter optimization phase involved a systematic evaluation of various neural network architectures and parameter settings to enhance model performance. The core model employed an attention-based transformer, which was iteratively trained and tested using different configurations. In the context of training neural networks, the process begins by feeding KING kinship data into the network, which then processes the data through a series of layers (in this case, both encoder and decoder layers) before producing an output. The error between the model’s predictions and the ground truth is computed and used to update the model’s parameters, aiming to minimize the error over time through backpropagation.

The model’s input consisted of a sequence of tokens representing individuals’ indices, with a maximum sequence length of 100, as determined by the network architecture. Transformer models project each input token into a fixed size embedding space and add positional embeddings of the same dimensionality, so the maximum sequence length is bounded by the size of the precomputed positional encodings and the number of learned positional vectors. The output was compared against the corresponding KING matrix values for the selected individuals using mean squared error (MSE) loss. Several key hyperparameters were explored during this phase, including the choice of activation function, the number of attention heads, and the number of layers in both the encoder and decoder. Additionally, the dimensionality of all layers was varied, and the learning rate for the ADAM optimizer was fine-tuned to ensure efficient convergence during training.

Note that our investigation was not of the performance ability of the model to predict the KING values, but rather of the model’s inner understanding of the individuals in relation to ecotype, or the heritage population of the individual.

### 4.3 Analysis

We assessed the quality of learned representations by analyzing embeddings extracted from the encoder layers of the attention transformer model before they were passed to the decoder. To facilitate interpretability, we applied UMAP directly to the KING matrix (Figure 8) and to the model’s learned embedding of the KING matrix (Figure 3) to project high-dimensional data into lower-dimensional spaces. This enabled visualization of population structure within these complex data for direct qualitative comparisons that informed quantitative assessments of QKV values in the latent space.

To further investigate model behavior, we utilized BERTViz, a visualization tool designed to analyze attention mechanisms in transformer models (Vig, 2019). Central to the self-attention mechanism are the QKV values — queries, keys, and values— which determine how attention weights are calculated and how information flows between tokens. By examining these QKV values, we gain insight into how the model prioritizes relationships and contextual dependencies within the genetic data. BERTViz enabled us to inspect the distribution of attention weights in the encoder layers, revealing not only the overall focus on relationships between individuals but also the underlying QKV dynamics that drive these patterns. All attention values shown in the heatmap figures are normalized to [0, 1]. This analysis provided critical human-readable insights into the interpretability of the model’s attention-based representations, reinforcing our understanding of how genetic similarities are captured and processed by the transformer architecture.

We further quantified the connectivity patterns between individuals by performing significance testing on the average intra-population and inter-population connectivity ratios, as derived from the QKV values across each attention layer. To assess these paired differences, we employed the Wilcoxon Signed-Rank Test (du Prel *et al*., 2010; Rosner *et al*., 2006). This non-parametric test is particularly well-suited for our analysis because it does not assume a normal distribution and effectively handles both ordinal data and continuous data with non-normal distributions. The Wilcoxon Signed-Rank Test evaluates whether the median difference between paired samples significantly deviates from zero, making it ideal for comparing our paired ratios of connectivity. Its robustness to outliers and skewed distributions ensures that our significance testing remains reliable, even when individual connectivity ratios vary widely or do not conform to standard parametric assumptions. We apply a log transformation to ratio data, a standard practice to stabilize variance and achieve normality (which is beneficial when performing non-parametric tests); we then compare the p-value to a significance level of *α* = 0.05, where our null hypothesis *H*_0_ is a ratio of 1 (no difference of ratios about the median).

## 5 Limitations

One of the main limitations of the Seq2KING model is the effort required to identify which encoder layers and attention heads display relevant and interpretable attention patterns. While the model learns meaningful representations of genetic relationships, the distribution of attention across multiple heads and layers varies, making it nontrivial to pinpoint the most informative regions. Extensive experimentation is often needed to determine which layers and heads contribute the most to capturing population structure and kinship relations. This issue is widespread as well in interpretability of LLMs used within industry; for example, analysis of BERT attention weights requires extensive testing of specific inputs and examination of layers (Clark *et al*., 2019).

Additionally, our model architecture imposes constraints on the number of individuals that can be processed simultaneously. Due to memory and computational limitations, Seq2KING can only handle sequences with a maximum length of 100 individuals at a time. However, the full dataset we tested consists of 2,503 individuals, preventing a direct assessment of connectivity across the entire population in a single pass. Instead, we rely on subsampling strategies to analyze different portions of the dataset, which may introduce variability in how relationships are captured across different runs. We note that from our tests of positional invariance on the contextual embeddings, we found that the embeddings of the same individual across different positions had very high cosine similarity; however, this invariance did not hold across different combinations of individuals.

The resolution of population labels used was at the continental level; the data contains the maximum resolution of origin country (and sometimes city), which we did not leverage in this study, thereby limiting our ability to validate the model’s performance at finer geographic scales and potentially masking more subtle sub-continental or local population structure. Future work could incorporate these higher-resolution labels to more rigorously evaluate Seq2KING’s capacity to detect micro-population differences. Doing so would also allow us to refine and calibrate attention-based interpretations against known demographic and geographic metadata.

We did not perform side-by-side comparisons with competing methods because these methods were not directly translatable to our framework. Existing approaches such as ADMIXTURE and fastmixture (Alexander *et al*., 2009; Santander *et al*., 2024) operate under fundamentally different assumptions and optimization constraints that are orthogonal to our work. ADMIXTURE estimates individual ancestry proportions by maximizing a likelihood model through block relaxation and quasi-Newton acceleration, thereby assigning individuals to a fixed number of ancestral populations. Similarly, Fastmixture builds on this framework by incorporating mini-batch expectation-maximization and randomized singular value decomposition (SVD) for initialization, which yields significant runtime improvements— up to 20–30*x* faster on large datasets. Both of these legacy population proportion-based methods rely on predefined population structure assumptions, modeling each individual’s genome as a mixture of fixed ancestral sources. In contrast, our transformer-based approach infers genetic relationships directly from kinship data without imposing explicit population labels, thus bypassing the need to predefine the number of ancestral clusters. As a result, direct benchmarking between our unsupervised, similarity-driven model and these proportion-based methods is not straightforward. Moreover, our goal was not prediction performance but understanding internal representations (see Methods).

Despite these current limitations, future iterations of our model will incorporate more rigorous benchmarking against alternative methods as we refine Seq2KING’s architecture. Active research is underway to develop evaluation metrics that allow for fair comparisons across different modeling approaches. With further improvements, we aim to establish Seq2KING as a competitive tool for analyzing genetic relationships and population structures in large-scale genomic datasets.

## 6 Conclusion

We further discuss this work in Supplemental Section 6.2, along with discussion of future work in Supplemental Section 6.2.1.

In this work, we introduced Seq2KING, a novel transformer-based model tailored for genetic relationship analysis. We began by leveraging data from the 1000 Genomes Project and processing it into a condensed KING matrix, which allowed us to capture pairwise relatedness coefficients and reduce the dimensionality of the genetic data. This approach addresses a fundamental computational challenge: a raw 30x coverage genome representation is approximately 50 gigabytes per human, making the representation of all humans (∼400,000 petabytes) computationally intractable. By using dimensionally compressed KING matrices, we theoretically enable the representation of relationships among all humans in a manageable format.

Our methods centered on neural excavating the QKV latent space - a challenging non-linear environment that offers the most granular information in artificial intelligence. Unlike legacy approaches that rely on discrete pre-defined labels, Seq2KING’s attention mechanisms provide continuous measures of relatedness, allowing us to theoretically visualize connectivity between any individual and all other humans. We found, in an unsupervised fashion, strong attention weight connectivity (85.34% intra-connectivity) within populations while maintaining structured, albeit weaker, connections across populations. To address the difficulty of visualizing these high-density attention weights, we introduced BERTViz to genetics, enabling human-readable analysis of these complex relationships.

The implications of our work extend beyond computational advances to applications in tracing migration patterns (such as Out-of-Africa journeys) and identifying subtle subgroupings within populations. By offering an unbiased method to explore population structure without pre-assigned labels, Seq2KING creates opportunities to remove background noise in genetic studies - a critical step for developing DNA-guided medications such as CRISPR therapies for genetic diseases affecting approximately 800 million people worldwide. Additionally, as an institutional model for latent space analysis, Seq2KING could serve as a contextual module within larger systems aimed at interpreting the human genome as a first language, potentially revealing the natural “family trees” hidden in our DNA.

## Supplementary material

### 6.1 S1 Code link

The code used to generate/format the data, the data itself, and all model-related code (training, inference, analysis, comparison) is publicly available on Github: https://github.com/EcotoneAI/Seq2KING

### 6.2 S2 Further discussion

We demonstrate in this work that the transformer model’s attention mechanism captures population structure by relating connections between genetically similar individuals while maintaining structured, albeit weaker, interactions across different populations. Similar to how words in a sentence influence one another based on contextual meaning, individuals in a population shape attention distributions through their genetic proximity. As we analyze attention patterns, we effectively trace a path through the dimension of similarity, observing how the model strengthens intrapopulation connections while preserving inter-population structure in a meaningful way. Our approach is theoretically effective on both small and large scales. On a small scale, for example in bacterial genomes, we anticipate being able to visualize pedigree trees that reveal forks of speciation and micro-evolutionary events. This is due to the model internalizing a form of population continuity, where attention strength is governed by genetic similarity, with attention flowing across layers, dynamically adjusting based on shared ancestry and relatedness. Moreover, when considering a large-scale scenario — such as constructing a pangenome for all eight billion humans — we envision generating a comprehensive relatedness map of humanity. This would enable unprecedented insights into global population structures and migrations over time.

A significant technical challenge in this work is interpreting patterns within the model’s latent space. Unlike explicit feature-based methods, transformers operate in high-dimensional representations that are not directly observable. This makes it difficult to extract clear rules governing how the model processes relationships between individuals. While we can visualize output embeddings or inspect attention distributions, these insights are inherently indirect. Deciphering what specific transformations occur within the query-key-value (QKV) mechanism remains a challenge (Pan *et al*., 2024), requiring tailored techniques for interpretability.

Despite these challenges, operating within the latent space of a transformer model is valuable because it provides the most granular and dynamic representation of relationships in current machine learning techniques (Marbut *et al*., 2024). The QKV mechanism allows the model to construct flexible and adaptive relationships between individuals, capturing nuances that fixed embedding methods might overlook. This makes it an ideal tool for modeling complex genetic structures, where relationships are neither strictly hierarchical nor purely distance-based. By examining the QKV values within the attention layers, we found strong connections in individuals within populations and weak connections in individuals across populations. These connections in the latent space were emergent and unsupervised as Seq2KING was not exposed to continental labels. This demonstrates the model capacity to learn population structure from the high dimensional manifold of the KING kinship matrix.

To manage the vast amount of information encoded in the transformer’s attention maps, we utilized the visualization tool BERTViz. This tool enabled us to display and interpret massive quantities of attention data, revealing intricate patterns that would be difficult to extract through numerical analysis alone. By leveraging BERTViz, we gain a more intuitive understanding of how the model distributes attention across individuals, offering a powerful means of inspecting and refining our approach. As transformer-based methods continue to evolve, these interpretability techniques will be crucial for unlocking deeper insights into the latent structures underlying genetic data.

#### 6.2.1 Future work

Various methods and tooling can expand and elaborate the work that we performed. For instance, our approach is translatable to population proportion frequency plots as found in ADMIXTURE-related papers (Alexander *et al*., 2009; Santander *et al*., 2024). This visualization depicts allelic frequencies of individuals across different populations and is valuable in both validating our current findings and new insights of population dynamics.

We also intend to incorporate PET files (chromosome-specific data files that contain pairwise kinship estimates, enabling detailed analysis of genetic heritage and relatedness at the chromosomal level), which will enable a more detailed kinship and heritage analysis by chromosome. The integration of chromosome-specific data will allow us to dissect genetic relationships with higher resolution and provide a clearer picture of inheritance patterns across the genome. Moreover, we will investigate alternative reduced forms of genetic representation. This includes exploring other versions or upgrades to the KING algorithm that have been proposed in recent literature. By examining these alternative approaches, we aim to determine whether enhanced versions of KING or entirely new methodologies can offer more precise or informative genetic relatedness coefficients.

Furthermore, we aim to develop methods that convert attention weight matrices directly into population or ecotype structures. By leveraging the detailed information captured in the self-attention mechanism, we aim to map these weights to explicit cluster assignments. This approach would allow us to bypass intermediate dimensionality reduction steps and directly extract population structures from the model’s learned representations. Such a method could facilitate more automated and precise cluster analysis, ultimately providing a powerful tool for uncovering hidden genetic substructures in large-scale datasets. We can extend our work by developing a cluster alignment graph, where identified continental population clusters can be easily compared to the ones found by performing UMAP on the original KING matrix. With the analytical tools established in this study, we will be able to identify and align clusters, providing a visual representation of how closely related individuals group together. This graph will enhance our understanding of population structure and facilitate further exploration of subtle genetic variations.

In addition to these initiatives, the integration of a diffusion transformer into our model could be integral. Diffusion transformers combine the denoising diffusion process with the powerful representation capabilities of transformers, enabling them to capture both global and local data structures effectively. In our case, adopting a diffusion transformer could enhance the model’s ability to handle the inherent noise and variability in genetic data, while also improving its sensitivity to subtle population differences. We anticipate that this approach will provide more robust embeddings and clearer insights into kinship patterns, further advancing our understanding of population dynamics.

Finally, the neural excavation tools developed in this paper lay the groundwork for future population discovery efforts. As these tools evolve, they will not only support more nuanced kinship and admixture analyses but also enable the discovery of previously unrecognized population substructures. We believe that these advancements will contribute significantly to the field of population genetics and open new avenues for research.

### 6.3 Supplementary tables

**Table S1:** Table of tested hyperparameters, that were varied across tests. All encoder/decoder layers composed of n number of attention heads, 2 linear feedforward layers, norm applied, and then given activation. MSE loss taken from validation at the end of all training epochs. The bolded row is the chosen architecture from which the embeddings were taken.

| Activation | Attention Heads | Encoder Layers | Decoder Layers | Dim of Linear Layers | ADAM Learning Rate | MSE Loss |
| --- | --- | --- | --- | --- | --- | --- |
| GELU | 1 | 5 | 5 | 3000 | 0.00001 | 0.00051 |
| GELU | 1 | 3 | 2 | 2000 | 0.00001 | 0.00051 |
| **Tanh** | **1** | **3** | **2** | **2000** | **0.00001** | **0.000031** |
| Tanh | 4 | 4 | 3 | 2000 | 0.0001 | 0.00039 |
| ReLU | 2 | 5 | 4 | 3000 | 0.0001 | 0.00039 |
| GELU | 2 | 5 | 4 | 3000 | 0.0001 | 0.00039 |
| Tanh | 2 | 5 | 4 | 3000 | 0.0001 | 0.0004 |

### 6.4 Supplementary figures

**Figure S1:**
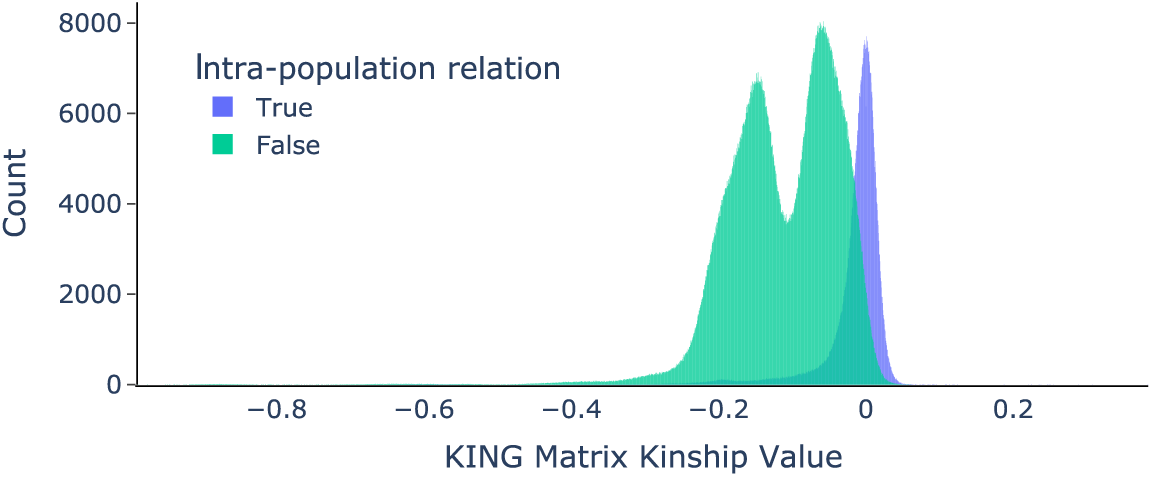
Histogram of KING Kinship values from the filled reflected matrix, colored by whether the value in the matrix is intra (is of a connection between individuals in the same population) or inter (connection between individuals in different populations).

**Figure S2:**
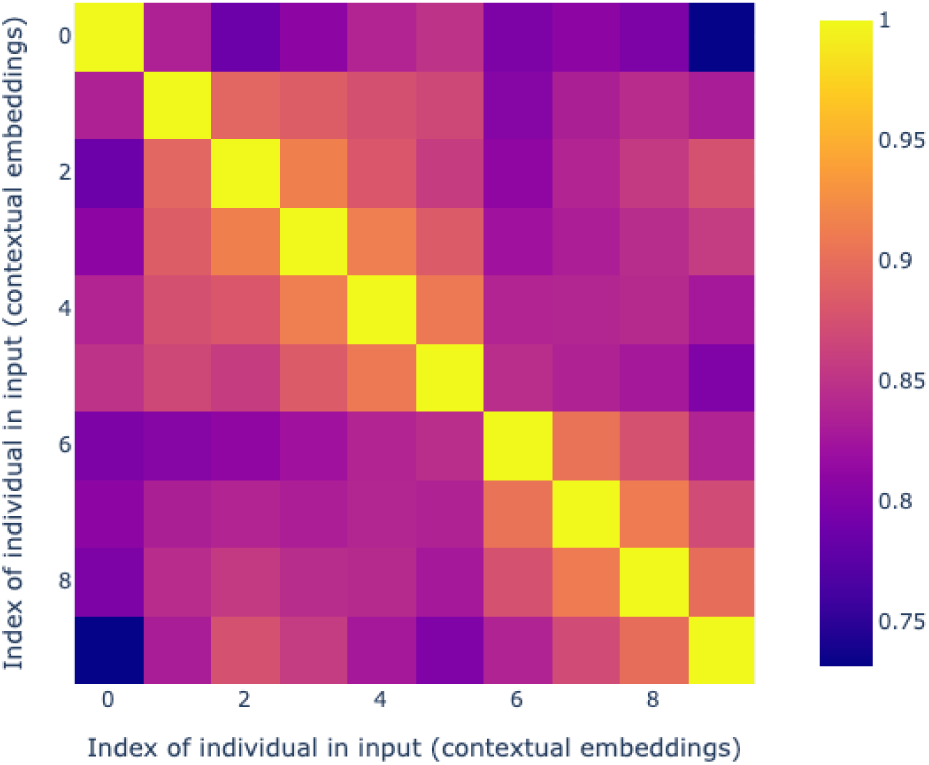
Cosine similarity matrix of the contextual embeddings of a given indivdual (id 42), acquired from moving the positional index of the individual from 0. . . 10 in a fixed sample input of 10 individuals. The other individuals stay fixed in their position in the input list. Tests positional invariance.

